# Alzheimer related genes show accelerated evolution

**DOI:** 10.1101/114108

**Authors:** Anne Nitsche, Kristin Reiche, Uwe Ueberham, Christian Arnold, Jörg Hackermüller, Friedemann Horn, Peter F. Stadler, Thomas Arendt

## Abstract

Alzheimer's disease (AD) is a neurodegenerative disorder of unknown cause with complex genetic and environmental traits. Here, we show that gene structures of loci, that show AD-associated changes in their expression, evolve faster than the genome at large. This phylogenetic trait of AD suggests a critical pathogenetic role of recent adaptive evolution of human brain and might have far reaching consequences with respect to the appropriateness of model systems and the development of disease-modifying strategies.

---

Alzheimer's disease (AD) is an age-related, chronic neurodegenerative disorder, neuropathologically characterized by fibrillar aggregates of A*β*-peptides and the microtubule-associated protein tau. While AD is extremely prevalent in human elderly, both A*β* and tau pathology are less common in non-primate mammals, and even non-human primates develop only an incomplete form of the disease [1]. This specificity of AD to human clearly implies a phylogenetic aspect of the disease and indicates that adaptive changes of cerebral structure and function that have occurred during human evolution may have rendered the human brain sensitive to AD [2]. Still, the evolutionary dimension of the AD pathomechanism remains difficult to prove and has not been established unequivocally so far.

To prove the contribution of brain evolution towards the AD pathomechanism, here, we established the AD-associated genome-wide RNA profile comprising both protein-coding (cRNA) and non-protein-coding (ncRNA) transcripts. For the first time, we also applied a systematic analysis on the conservation of splice sites based on multiple alignments [3] across 18 vertebrates of homologs of AD-associated protein-coding and noncoding genes [Supplement Sec. 1.8].

To this end, we designed a custom array comprising 931,898 probes derived from Agilents Whole Human Genome Oligo array, long non-coding RNA (lncRNA) probes extracted from public databases, computationally predicted loci of structured RNAs, and lncRNA probes experimentally identified by transcriptome-wide expression variation studies based on the Affymetrix Human Tiling 1.0 array comparing AD patients with control samples [Supplementary Methods]. Applying this custom array to 19 AD patients and 22 control samples, we identified a differential expression of 154 multi-exonic cRNAs with a total of 4,162 splice sites and 141 multi-exonic lncRNAs with a total of 1,297 splice sites.

Genome-wide studies that systematically analyze the evolutionary age of protein-coding and non-proteincoding AD-associated genes have not been performed previously. While major evolutionary changes might have occurred at the transcriptomic level, they appear to be particularly pronounced for lncRNAs [3, 4]. As shown by analyses of sequenced genomes of a large variety of species, the relative amount of non-coding sequence increases consistently with complexity [5]. Thus, lncRNAs, most likely constitute a critical layer of gene regulation in complex organisms that has expanded during evolution [6]. However, the evolutionary histories of lncRNAs have been notoriously hard to study due to their usually low level of sequence conservation. This not only hampers comprehensive homology-based annotation efforts but also makes it nearly impossible to obtain the high fidelity sequence alignments that are required for in depth studies into their evolution. Alternatively, the conservation of gene structure and particularly the conservation of splice sites may also be used to establish homology of lncRNAs [3]. Splice sites therefore leave “phylogenetic footprints”, and conserved patterns of splice sites may be used to predict novel transcripts from multiple genome alignments [7, 8]. Although lncRNAs are clearly ancient components of vertebrate genomes, they exhibit a rapid turnover of their intron/exon structures [3] that may be indicative of functional adaptation.

While the disease-relevance of lncRNAs is increasingly recognized, previous systematic gene expression profiling studies in AD nevertheless focussed predominantly on protein-coding genes. Consequently, so far, only a few individual AD-associated ncRNAs have been identified and functionally characterized [9].

In order to compare the conservation of genes at a structural level, we classify the data by the “degree of conservation” *c*, which is the fraction of conserved splice junctions per gene. We ask – for a fixed value of *c* – whether loci that are differentially expressed in AD patients show signs of accelerated evolution compared to the set of genes contained in the Gencode v14 annotation of the human genome. Different thresholds of c highlight different aspects of conservation and evolutionary change: At *c* > 0% we assay only presence or absence of a gene, and thus its evolutionary origin. The other extreme, *c* = 100% focusses on the precise conservation of the gene structure. Protein-coding genes and non-protein-coding genes were independently investigated for their conservation [Supplement, Fig. S1].

Nearly all AD-associated protein-coding genes are evolutionarily old (Fig. 1C). There were no differences in conservation rate at *c* > 0% between AD-associated and all protein-coding genes, i.e., AD-associated protein-coding genes did not originate later in evolution than other protein-coding genes. In line with previous reports [6], lncRNAs are much less well conserved and many have emerged in the course of mammalian evolution. The fraction of conserved lncRNAs thus decreases rapidly with evolutionary time (Fig. 1A,B). As for protein-coding sequences we did not observe a significantly younger origin of AD-associated genes.

**Figure 1:**
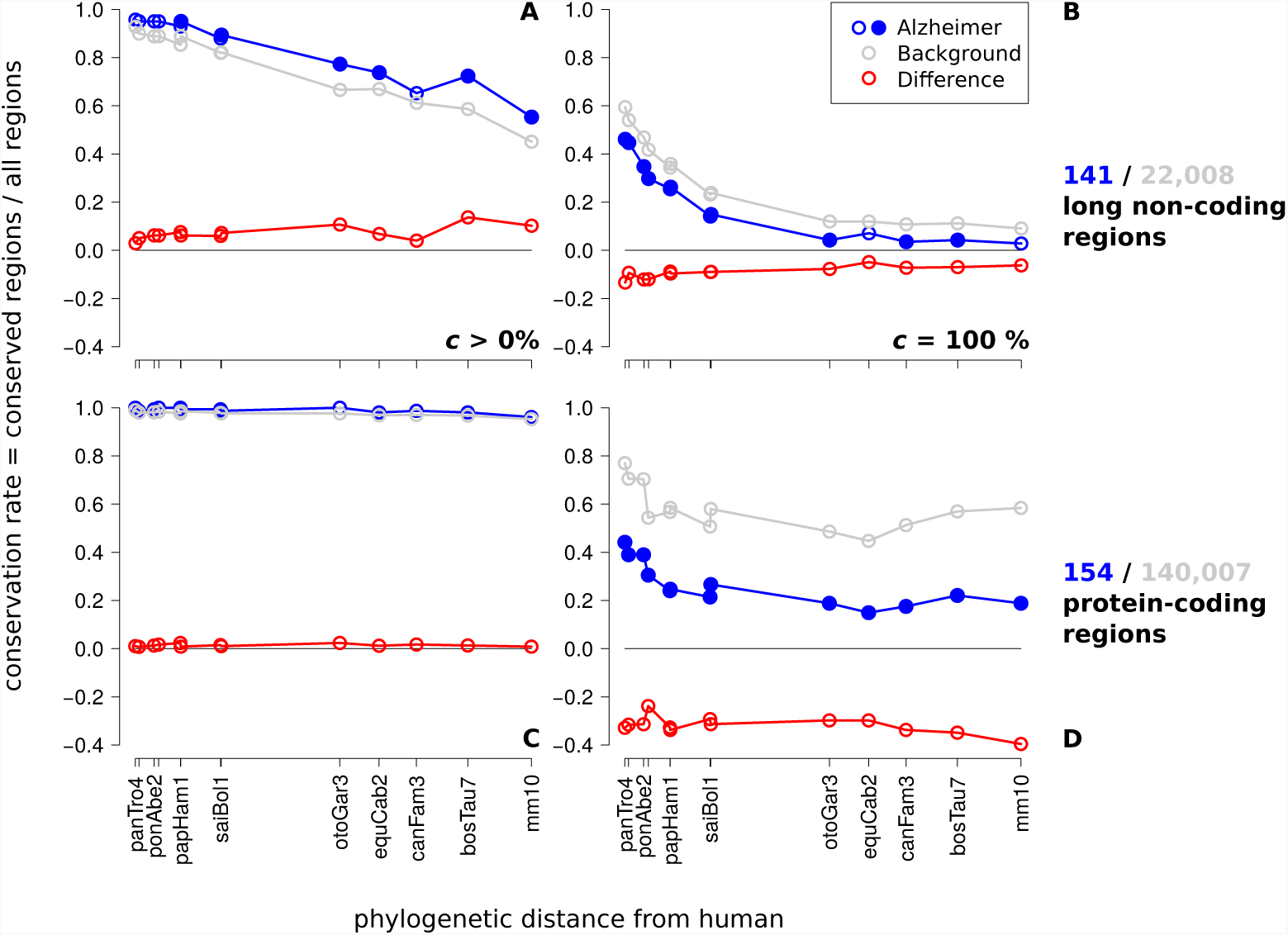
Conservation rates of human AD-associated non-protein-coding (A,B) and protein-coding (C,D) regions. Comparing presence (*c* > 0%) and precisely conserved gene structure (*c* = 100%). On the horizontal axis mammalian species are indicated (denoted by the UCSC abbreviations) at their phylogenetic distance from human. Distinct data points are connected by lines to guide the eye. Variations in assembly and alignment quality cause some non-monotonicity in the curves, the overall decrease of conservation with phylogenetic distance is nevertheless clearly visible. Statistical significance of differences is computed independently for each species. Filled circles indicate *p* < 0.05. The fraction of detectable conserved AD-associated non-coding genes is marginally higher than the conservation of the background set non-coding transcripts if only presence/absence of a transcript is considered (**A**). In contrast, if conservation of the entire gene structure is considered, AD-associated genes are significantly less conserved than the control. This is true for both lncRNAs (**B**) and protein-coding genes (**D**). The Supplement provides further conservation rate results for conservation degree *c* > 60%. Additional controls against possible confounding effects e.g. of alignment quality in Supplement Fig. S4 and Fig. S5 show that the trends shown here are robust.

While there is no recognizable difference in the evolutionary age of origin between AD-associated genes compared to the transcriptome as a whole (Fig. 1C), we observed significant, albeit more subtle differences in the evolution of AD-associated and general lncRNAs, concerning the changes in gene structure. With an increasing degree of conservation c, the conservation rate of AD-associated non-coding genes decreases and becomes significantly (*p* < 0.05) distinguishable from the background level (Fig. 1A,B) not only for distantly related mammals but even primates, when complete conservation of gene structure is considered (*c* = 100%). In other words, the fraction of transcripts that have the entirety of their splice sites conserved is smaller amongst AD-associated ncRNAs than amongst non-coding genes at large. AD-associated ncRNAs hence show an accelerated evolution of their gene structure. This is indicative of a more rapid functional adaptation of AD-associated non-coding genes.

Although protein-coding genes are much better conserved than lncRNAs we observed the same increase of splice site turnover in AD at *c* = 100%. In fact, the relative effect is even stronger compared to nonprotein-coding loci (≈ 30-40% *versus* ≈ 5-15% difference, shown as red lines in Fig. 1B and 1D, respectively). Since the same fraction of transcripts is already detectable at low conservation degrees, while the conservation rate decreases with higher *c*, we conclude that splice sites are systematically less conserved in human AD-associated regions compared to the typical behavior of the transcriptome.

While protein-coding loci exhibit an enhanced rate of small changes in their gene structure, we observe large changes in lncRNAs, again with a significantly enhanced rate in the AD-associated ncRNAs. This suggests that in particular AD-associated non-coding genes play an important, as yet largely unexplored, role in the AD pathomechanisms.

We have shown here that gene structures of AD-associated loci evolve faster than the genome at large, while there is no evidence that AD-associated genes originated particularly late in evolution. In order to capture the evolution of lncRNAs, we focused on gene structure, i.e., the conservation of splice sites because this approach makes it possible to separate the evolution of the transcripts from other selective constraints such as regulatory DNA elements that may affect sequence conservation [3]. Changes in gene structure can be expected to have in general larger functional effects than point mutations. The enhanced rate of gene structure evolution in AD-related genes hints a relation of AD to recent adaptive evolution, presumably in relation to the rapid evolution of the human brain. Importantly, replacing the background set by only gene expressed in brain did not affect the conclusions [Supplement, Sec. 2.3].

Major phenotypic brain changes that have occurred in the course of recent human evolution, in particular between human and chimpanzee, appear to be mostly the result of an increase in gene expression and are, thus, reflected at the transcriptomic level. [10–12]. Genes whose expression has increased in human brain are mainly related to growth and differentiation [13] and frequently are involved in transcriptional regulation and RNA processing [10, 11]. The most significant differences in gene expression between the human and nonhuman primate brain have been observed in the association cortex [10, 14], i.e., brain areas that have expanded during hominid evolution [15] and are affected in AD most early and most constantly [16]. Evolutionary expansion of the neocortex, and in particular phylogenetic shaping of association areas, is associated with a developmental deceleration and an extended period of high neuronal plasticity into adulthood [13]. The presence of these neurons which remain structurally immature throughout their lifespans might provide the prerequisite both for the human adaption to the “cognitive niche” and for a high vulnerability towards factors that lead to the development of AD [17–19].

Our data support the concept that neuronal vulnerability in AD is a result of the evolutionary legacies that have occurred during the course of evolution of the human brain, making AD an example of antagonistic pleiotropy. This evidence for a phylogenetic trait of AD highlights the necessity to reconsider our approaches to define the molecular pathology of AD and the appropriateness of current animal model systems [20] to develop disease-modifying strategies.

## Acknowledgements

We are indebted to Stephan Schreiber for expert technical assistance. This work was supported in part by DFG grants within SPP1738 (AR 200/17-1, HA 5857/2-1) to PFS, TA and JH.

## Author contributions

TA, UU, KR, CA, and JH designed and performed the tiling array and custom array experiments. KR, CA, and JH performed bioinformatic analysis of the expression data. AN developed and performed the computational method to assess the conservation of splice sites.

